# Evolution of *Drosophila buzzatii* wings: Modular genetic organization, sex-biased integrative selection and intralocus sexual conflict

**DOI:** 10.1101/2021.06.16.448721

**Authors:** PP Iglesias, FA Machado, S Llanes, E Hasson, EM Soto

**Affiliations:** Laboratorio de Genética Evolutiva, Universidad Nacional de Misiones – CONICET, Félix de Azara 1552, N3300LQH, Misiones, Argentina; Department of Biological Sciences, Virginia Polytechnic Institute and State University, Blacksburg, United States; Instituto de Ecología, Genética y Evolución de Buenos Aires (IEGEBA – CONICET), DEGE, Facultad de Ciencias Exactas y Naturales, Universidad de Buenos Aires, Buenos Aires, Argentina

**Keywords:** Adaptive landscapes, Genetic architecture, Intralocus sexual conflict, Morphological integration/modularity, Multivariate selection

## Abstract

The *Drosophila* wing is a structure shared by males and females with the main function of flight. However, in males, wings are also used to produce songs, or visual displays during courtship. Thus, observed changes in wing phenotype depend on the interaction between sex-specific selective pressures and the genetic and ontogenetic restrictions imposed by a common genetic architecture. Here, we investigate these issues by studying how the wing has evolved in twelve populations of *Drosophila buzzatii* raised in common-garden conditions and using an isofemale line design. The between-population divergence shows that sexual dimorphism is greater when sex evolves in different directions. Multivariate Qst-Fst analyses confirm that male wing shape is the target for multiple selective pressures, leading males’ wings to diverge more than females’ wings. While the wing blade and the wing base appear to be valid modules at the genetic (**G** matrix) and among-population (**D** matrix) levels, the reconstruction of between-population adaptive landscapes (**Ω** matrix) shows selection as an integrative force. Also, cross-sex covariances reduced the predicted response to selection in the direction of the extant sexual dimorphism, suggesting that selection had to be intensified in order to circumvent the limitations imposed by **G**. However, such intensity of selection was not able to break the modularity pattern of the wing. The results obtained here show that the evolution of *D. buzzatii* wing shape is the product of a complex interplay between ontogenetic constraints and conflicting sexual and natural selections.

## Introduction

Understanding changes in morphological structures requires an integrative approach that also considers constraints upon change. How is morphology produced during development in the first place? Is selection in line with these developmental rules? Does selection differ between the sexes? Morphological integration, selection, and between-sex pleiotropy are key factors generating association among traits at larger scales. An integrated developmental pattern or the occurrence of sex-specific selection on homologous structures controlled by common genetic machinery (i.e. intralocus sexual conflict) may constrain the evolutionary trajectories of such structures. It is only after evaluating the relative effect of these factors that morphological changes can be understood in the light of the interaction between constraints and their potential for functional adaptation.

The wing of *Drosophila* is a complex structure involved in different functions such as flight and acoustic or visual communication (Wooton, 1992). For a long time, it has been considered as a developmentally integrated structure that constrains adaptive evolution (Houle, Bolstad, Van der Linde, & Hansen, 2017; Klingenberg, 2009; Klingenberg & Zaklan, 2000). While these conclusions are mostly based on the hypothesis that the wing is divided into anterior and posterior (AP) compartments (Klingenberg & Zaklan, 2000), recently Muñoz-Muñoz et al. (2016) showed evidence supporting the compartmentalization of this structure along the proximo-distal (PD) axis, forming two different modules, the wing base and the wing blade. These modules were recognized not only on the phenotypic level but also at the genetic and environmental levels, suggesting that it is possibly a consequence of a modular developmental program.

From the wing primary task perspective (i.e. flight), the developmental modules seem to match functional modules: the wing base transmits the forces generated by the flight muscles and the wing blade generates the aerodynamic forces necessary to lift the body (Dudley 2002). Although fly’s ability is common to both sexes, sex-biased dispersal has been documented in *Drosophila* (Begon, 1976; Powell, Dobzhansky, Hook, & Wistrand, 1976; Fontdevila & Carson, 1978; Markow & Castrezana, 2000; Mishra, Tung, Shree Sruti, Srivathsa, & Dey, 2020). An asymmetric dispersal of the sexes can exert different selective pressures on wing morphology in each sex, leading to a sex-biased evolution of the modules.

*Drosophila*’s wings are also involved in premating behaviors that markedly differentiate the wing’s function between sexes (Ewing, 1983; Dickson, 2008). Except for a few species (within the *Drosophila virilis* species group; Satokangas, Liimatainen, & Hoikkala, 1994), only males use wings for acoustic or visual communication. Therefore, if wing morphology influences sound production or visual display, only male wings will be subject to selection. This selection on males can cause the displacement of females from their phenotypic optimum, reducing their fitness. It is well documented that the direction and intensity of selection on courtship song have been found to differ among populations and species according to female preferences (Iglesias & Hasson, 2017; Iglesias et al., 2018a; Klappert, Mazzi, Hoikkala, & Ritchie, 2007).

Here, we address these issues by investigating how the wing has evolved in twelve populations of the cactophilic species *Drosophila buzzatii* raised in common-garden conditions and using an isofemale line design. Only the males of the *D. buzzatii* species use wings to produce a courtship song (Iglesias & Hasson, 2017; Iglesias et al., 2018a; Iglesias, Soto, Soto, Colines, & Hasson, 2018b), and a previous study has shown a rapid divergence of courtship song parameters among these populations (Iglesias et al., 2018a). Thus, we first investigated whether the wings of males and females are evolving differently among the sampled populations. At present, wing modularity at the genetic level has only been tested in the model species *D. melanogaster* (but see Soto, Carreira, Soto, & Hasson, 2008); however, *D. buzzatii* is a cactophilic species distantly related to it. The effect of selection and drift accumulate through time in each evolving lineage and can even change the modular pattern in an evolutionary context (Martín-Serra, Figueirido, & Palmqvist, 2020; Melo & Marroig, 2015). Thus, we then tested whether the wing in each sex is organized in two modules along the PD (proximo-distal) or the AP (antero-posterior) axis in *D. buzzatii*. We test these hypotheses by estimating the posterior distribution of the Among-Population (**D**) and Additive Genetic (**G**) within and between sexes covariances from a landmark-based analysis. We also estimated the covariance pattern among peaks on the realized adaptive landscape for these populations (**Ω)** to evaluate the potential effect of selection in defining between-populations patterns of phenotypic integration and modularity. We also applied a multivariate ***FSTq*** *–FST* approach to determine the extent to which selection and drift contributed to the evolution of the wing. Finally, we investigated how cross-sex covariances constrain or facilitate the predicted responses to multivariate selection. To do that, we combined the multivariate breeder’s equation with random and empirical selection gradients and compared the predicted response to selection when using a **G** with and without between sex covariances. If intralocus sexual conflict is not fully resolved, we expect high and positive intersexual genetic correlations resulting in an augmented response when cross-sex covariances are setting to zeroes. Otherwise, if the genetic architecture of wings evolves to allow independent adaptation in each sex, thus resolving intralocus conflict, we expect the response to selection to be the same. However, if cross-sex covariances facilitate the response to selection, we expect the magnitude of the response to be higher when including them.

## Materials and Methods

### Data collection and measurements

For this study, we analyzed 12 populations of *D. buzzatii* flies that have been previously used to study variation in male courtship song (Iglesias et al., 2018a). Each population was characterized by eight to 15 isofemale lines founded with wild-collected gravid females. Flies were raised under common-garden and controlled-density conditions (40 first-instar larvae per vial), and a photoperiod regimen of 12-h light: 12-h dark cycle. First, they were raised on standard *Drosophila* medium for four generations and then were moved one more generation to a ‘semi-natural’ medium prepared with fresh cladodes of the cactus *Opuntia ficus indica* (see Iglesias et al., 2018a for more details). This cactus species represents the more widespread host used by *D. buzzatii* in the study area.

We removed the right-wing of between three to five adult emerged flies per sex and line, and mounted them on glass microscope slides for image acquisition. Following Muñoz-Muñoz et al. (2016), a set of 15 landmarks was digitized in each wing using the TPSdig software (Rohlf, 2001). Shape information was obtained from the configurations of landmarks using standard geometric morphometrics methods as implemented in the *Geomorph* package v3.0.7 (Adams, Collyer, Kaliontzopoulou, & Sherratt, 2018). To examine shape variation, we performed a principal component (PC) analysis based on the covariance matrix of Procrustes residuals. To evaluate how many PCs should we retain as wing shape variables in subsequent analyses we employed the broken-stick model criterion implemented in the package vegan (Oksanen et al., 2019), which indicated that only the first 5 axes should be kept (68.28% of total variation).

### Genetic covariation, population divergence and selection

We estimated the posterior distribution of the Among-Population (**D**) and additive genetic (**G**) covariance matrices by using a Bayesian multivariate mixed model implemented in the R package *MCMCglmm* (Hadfield, 2019). Population and Line (nested within Population) were included as fixed and random factors, respectively. The analysis was run for 10^5^ generations with 50% burn-in, after which we extracted 500 samples. Convergence was verified using trace plots for all parameters.

The covariance matrices for the fixed effect Population and random effect Line were considered as the Among-Population (**D**) and the genetic (**G**) covariance matrices, respectively. Each sex-trait combination was treated as a different trait (Sztepanacz & Houle, 2019) resulting in 10 traits for each level of comparison. Thus, **G** and **D** are composed of four submatrices representing both patterns of integration and divergence between and within sexes as follows

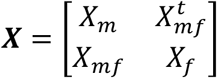

where ***X*** is the full covariance matrix, ***X***_***m***_ and ***X***_*f*_ are the male and female specific covariance submatrices, and *X*_*mf*_ and 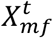 are the between-sex covariance submatrices, with *t* denoting a transpose. For **G**, the *X*_*mf*_ submatrix is also called **B**, and encodes the pleiotropic trait association and constraint between sexes.

To evaluate the potential effect of selection in defining between-populations patterns of divergence, we calculated the **Ω** matrix, which expresses the covariance pattern among peaks on the realized adaptive landscape for these populations. **Ω** is calculated as Ω = *G*^−1^*DG*^−1^(Felsenstein, 1988; Marroig, & Cheverud, 2010), which assumes that all populations share a single and unique common ancestral population. Ω was calculated for each sample of the posterior distribution, producing a total of 500 matrices.

Because all matrices were calculated on a reduced morphospace comprised of only five PCs, for each sample of the posterior we back-projected all matrices on the original image space to obtain the landmark variation associate with **G, D** and **Ω**. The covariance matrix **Z** on the original space can be obtained as follows *Z* = *VXV*^*t*^ where *X* is either **G, D** or **Ω** calculated on the first five PCs (as above), **V** is a matrix of the leading five eigenvectors. These back-projected matrices were then used on our modularity analysis.

### Sexual dimorphism in the sampled populations

To determine whether wing’s shape changes are sex-specific, we calculated the magnitude (Procrustes distance) and alignment (vector correlation) of between-sex shape differences for each sample of the posterior distribution of the fixed effect Population. For each population, the magnitude of the average differentiation between males and females in Procrustes distance was plotted against the Fisher z transformed vector correlation between males and females. A positive correlation between these variables is indicative that between sexes differences are accumulating along the same lines of divergence. A negative correlation would indicate that sexes tend to diverge in opposite directions, independently from each other.

### Modularity analyses

Muñoz-Muñoz et al. (2016) found evidence supporting the hypothesis of wing modularity along the PD axis in *D. melanogaster*. However, locating the boundaries between the modules is hard given that development is principally characterized by variations in terms of gradients (Matamoro-Vidal, Salazar-Ciudad, & Houle, 2015). Muñoz-Muñoz et al. (2016) also discuss the role of the wing hinge contraction during late pupal development as an important process potentially impacting the covariance structure and thereby on the PD modularity. However, given that hinge contraction occurs while cells at the margin of the blade are attached to the surrounding pupal cuticle (see Figure 5 in Matamoro-Vidal et al., 2015), it is possible that the proximal portion of the blade covariate more with the hinge than with the distal portion of the blade. Thus, whether to include landmarks 8 and 9 (following Muñoz-Muñoz et al., 2016) in one module or another is somewhat arbitrary. Therefore, in this study, we tested two hypotheses for the PD axis that only vary in assigning those landmarks to either the blade or the base. Hypothesis A of PD modularity includes landmarks 8 and 9 in the wing base module (Figure 1A) while hypothesis B includes those landmarks in the wing blade module as Muñoz-Muñoz et al. (2016; Figure 1B). For comparative purposes, we also tested the classical hypothesis in which the wing is divided into an anterior and a posterior compartment following Klingenberg (2009; also see Klingenberg & Zaklan, 2000; Figure 1C).

**Figure 1.**
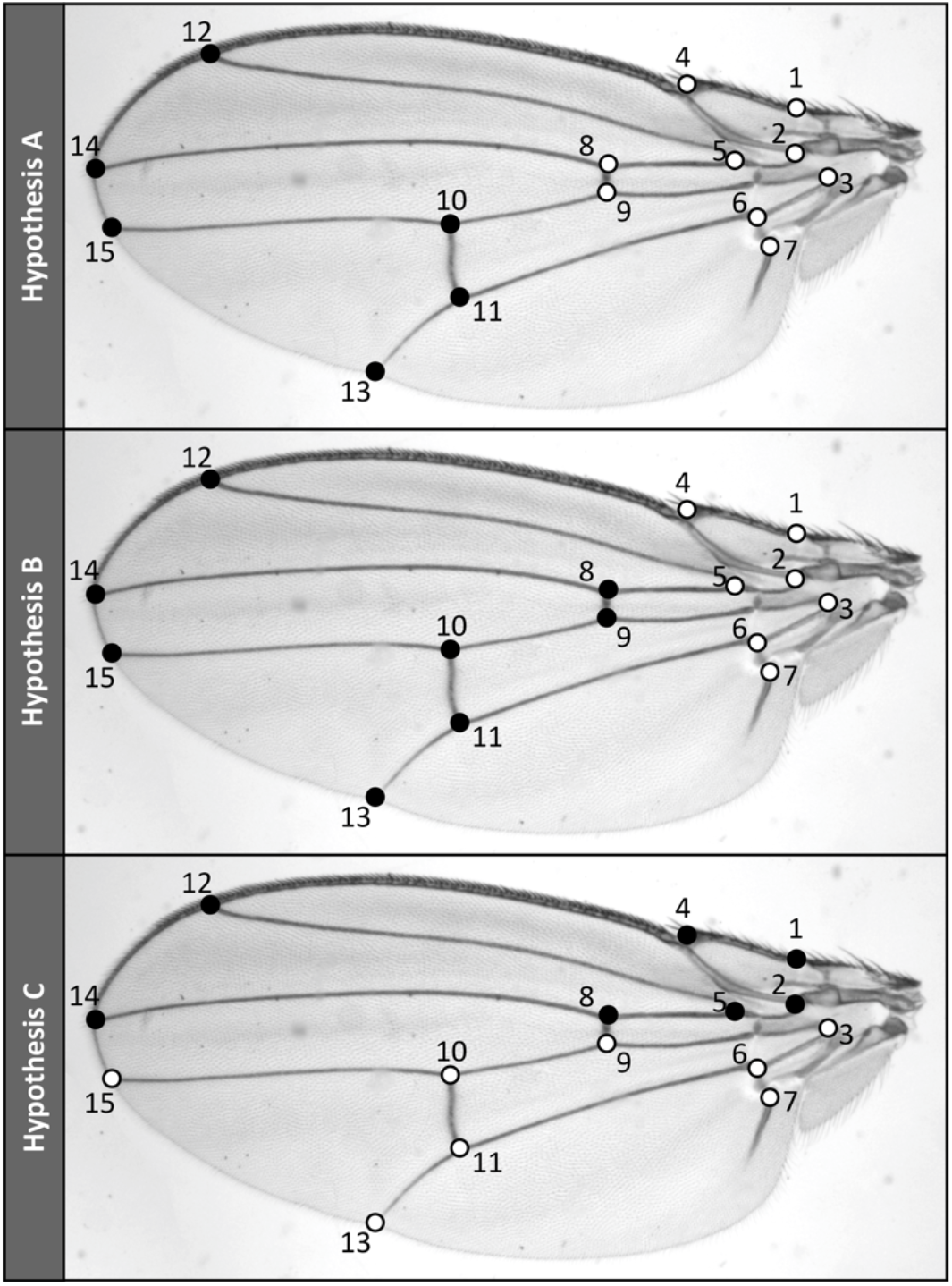
Digitized landmarks showing the three hypotheses tested for wing modularity. Landmarks of the same color belong to the same module.

The hypotheses of modular organization of the wing were evaluated using the covariance ratio effect sizes (Z_CR_) statistic (Adams & Collyer, 2019), which was shown to be robust even for small sample sizes. Z_CR_ is a function of the covariance ratio statistic (CR) which measures the intensity of association between modules in relation to the intensity of trait association within a module as measured by the covariance among those traits (Adams, 2016). Z_CR_ is then calculated as a transformation of the observed CR value (CR_obs_) as follows

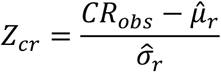

were 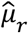 and 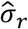 are, respectively, the average and standard deviation of CR under the null distribution of no modularity, and are calculated by shuffling matrices rows and columns 1000 times. For each matrix type, Z_CR_ was calculated for each sample of the posterior, producing 1000 values for each matrix. More negative values of Z_CR_ indicates a more modular structure, and positive values indicate less modular structures. Because Z_CR_ was proposed to compare the modularity signal among different datasets, here we use this statistic to both select the best modularity hypothesis for **G** (Figure 1), and to compare modularity signal among **G, D** and **Ω**. For **G**, the best hypothesis was chosen as the one with the lower Z_CR_. The best genetic hypothesis was then used to calculate modularity signals on the other levels of analysis. For all matrices, Z_CR_ was calculated for each MCMC sample, resulting in a posterior distribution of modularity signal values.

### Multivariate *FSTq –F*ST comparisons

To evaluate if phenotypes were evolving due to selection or drift we employed Chenoweth & Blows (2008) multivariate generalization of the QST-FST test called ***FSTq****–FST*. ***FSTq*** is a matricial transformation analogous to the calculation of the univariate Qst and can be obtained as follows:

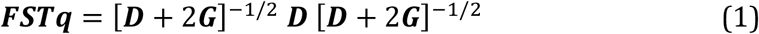

Under neutrality, the *i*th eigenvalue of ***FSTq*** (λ_i_) is expected to be equal to *FST* estimates. Eigenvalues that are larger than *FST* are thought to be generated by directional selection, and values that are lower are thought to be generated by stabilizing selection (Chenoweth & Blows, 2008). In this way, we can evaluate how directions of divergence were differentially affected by selection.

One common issue in multivariate systems is the overestimation of the leading eigenvalues due to sampling error (Marroig et al., 2012). Thus, using *FST* as the sole parameter as the null-expectation for λ_i_ can be considered unrealistic. Here we circumvent this issue by producing a distribution of expected eigenvalues using simulations, and confronting them against the observed ones. This was done by obtaining the expected patterns of between-population divergence E(**D**) as follows

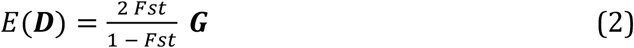

(Martin, Chapuis, & Goudet, 2008) using the **G**s from the posterior sample as a starting point. To simulate population divergence and to account for sampling error, we generated 12 observations for each E(**D**) and calculated the covariance among those values. This new covariance matrix of divergence was then used in equation (1) to calculate an empirical distribution of ***FSTq*** under drift. This was performed 10000 times by randomly drawing Gs from the posterior. Each simulated ***FSTq*** was subjected to an eigendecomposition and their eigenvalues were used as a null expectation for each dimension of the empirical ***FSTq***. The null distribution was then confronted against the observed values, both for individual axes (λ_i_) as well as for the average eigenvalue 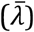 as a global test of the adaptive hypothesis. This analysis was performed for the full **G** and for **G**_**f**_ and **G**_**m**_ separately. Additionally, we computed the amount of between population divergence for each sample of the posterior distribution of ***FSTq*** as the sum of the variances for males and females specific traits individually. This produces a measure of how much each sex diverged from the ancestral population. We also conducted QST–FST comparisons using the R package *driftsel* (Karhunen, Merilä, Leinonen, Cano, & Ovaskainen, 2013).

The coancestry coefficient (*FST*) for each pair of populations were estimated by using data from eight microsatellite markers previously genotyped (Iglesias et al., 2018a) and the bayesian R package *RAFM* (Karhunen, 2012). Analyses were run for 10^6^ generations with a 50% burn-in, resulting in 500 posterior distribution samples.

### Cross-sex (co)variances and the response to selection

Finally, we investigated how cross-sex genetic covariances may constrain or facilitate the response to selection combining the multivariate breeder’s equation, the random skewers method and the R metric (Sztepanacz & Houle, 2019). The multivariate breeder’s equation (Lande, 1979) can be written to includes differences in selection and inheritance in the two sexes as follows:

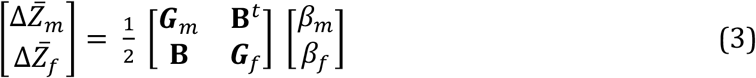

where Δ*<u>Z</u>* and *β* are the selection responses and selection gradients, respectively, in males (m) and females (f). By generating a large number of random selection gradients and applying to our **G** estimates we can evaluate how those populations will evolve under natural selection (Cheverud & Marroig, 2007). To investigate the constraining effect of between-sexes covariances (**B**), one can confront the magnitude of evolution (the norm of 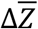) between a **G** with and without between sex covariances (**B** is set to a matrix of zeroes; Sztepanacz & Houle, 2019). If the ratio between both magnitudes (the R metric) is less than one, then the cross-sex genetic covariances can constraining the response to selection, whereas R>1 suggests they are facilitating the response to selection. We calculated R by drawing a vector from a spherical multivariate normal distribution and applying it to a randomly drawn G matrix from our posterior distribution. This was performed 10000 times, producing a distribution of Rs. Complementarily, we also calculated the R statistic for the empirical divergence between populations. We did that by rearranging equation 3 to estimate *β*s from 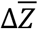 drawn from our posterior distribution of fixed effects. We then used these empirically derived *β*s to calculate a distribution of empirical R values as above.

## Results

### Sex-specific wing shape in the sampled populations

The inspection of the relation between magnitude (Procrustes distance) and alignment (z-transformed vector correlation) of between sex divergence for each of the 12 populations of *D. buzzatii* show a negative association (Figure 2A). The distribution of correlation among alignment and magnitude showed a 95% highest density interval that was entirely negative, with a median correlation of -0.629 (Figure 2B). This result suggests that higher sexual dimorphism was achieved when males and females evolved in different directions instead of in the same direction.

**Figure 2.**
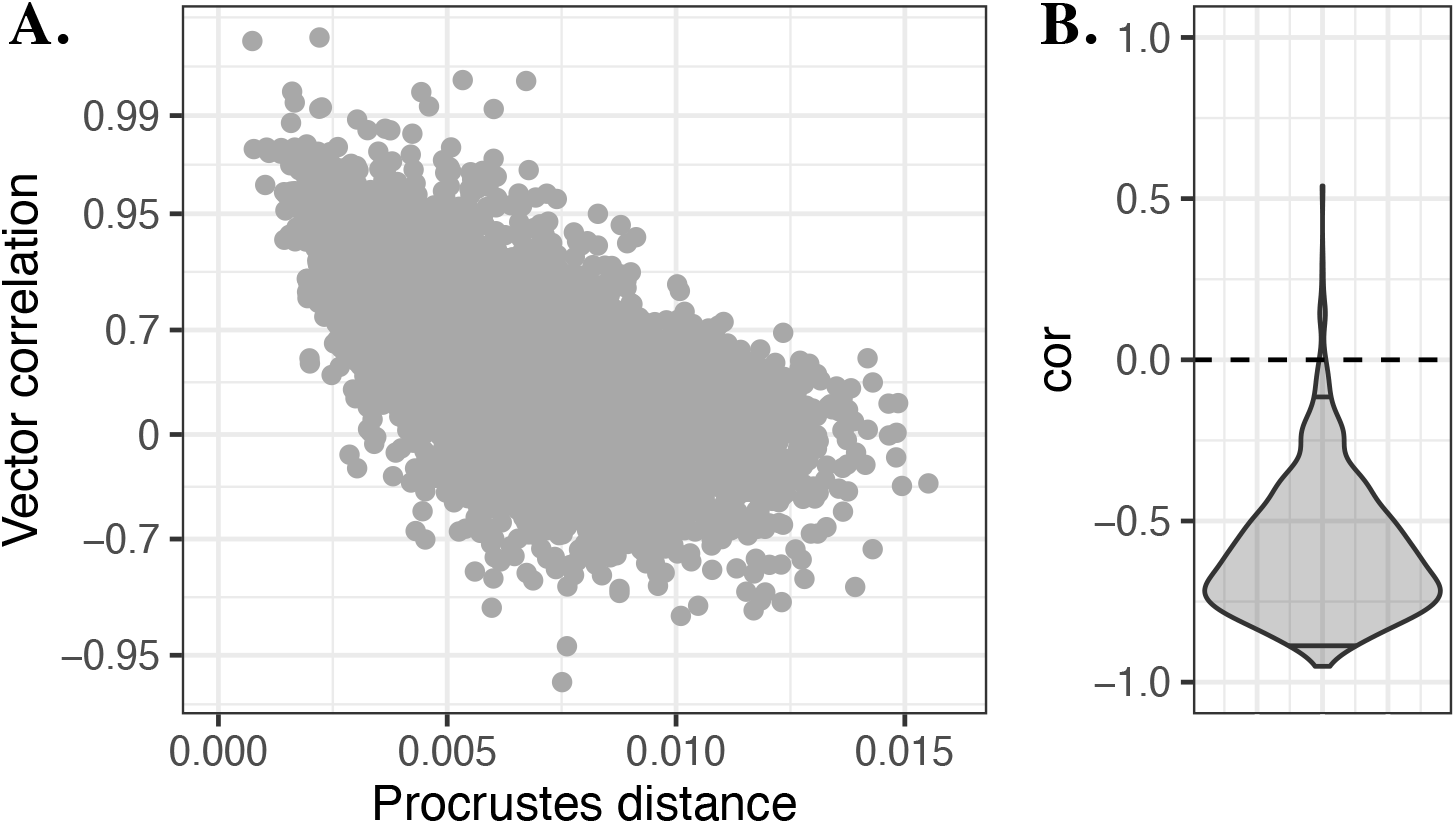
Magnitude (Procrustes distance) and alignment (vector correlation) of between-sexes shape differentiation among the 12 populations of *D. buzzatii*. **A**. Representation of between-sex divergence and alignment obtained from the posterior distribution of fixed effects. The Y axis is shown in a fisher-z transformed scale. **B**. Distribution of Pearson-correlation values between Procrustes distance and fisher-z transformed vector correlation for all samples of the posterior distribution. Horizontal solid lines inside the distribution highlights the 95% higher density intervals.

### Patterns of phenotypic integration and modularity

The analysis of the modularity signal through Z_CR_ on **G** suggests that our PD hypothesis A (H_A_) is the best one for both males and females (Figure 3A). For H_A_, all Z_CR_ from the posterior distribution were significantly different from 0 for both sexes (p<0.001), with values being generally lower than -2. While the hypothesis B (H_B_) also showed values that were generally significant, the intensity of modularity signal was weaker, with values closer to zero (no modularity). Lastly, the classical antero-posterior hypothesis C (H_C_) showed values that were consistently non-significant (p>0.10) and were larger than zero.

**Figure 3.**
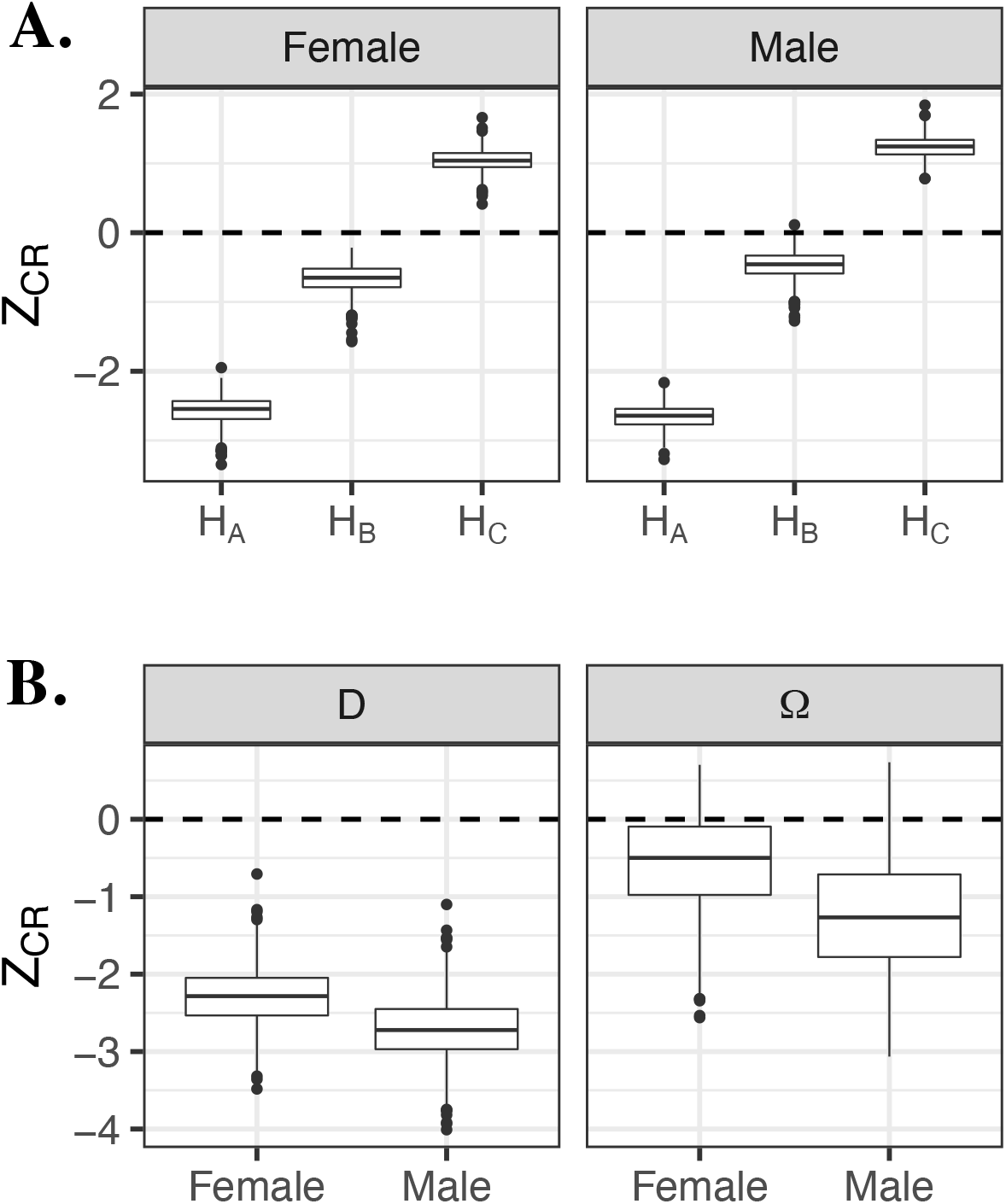
A. The posterior distributions of the Z_CR_ statistic for modular hypothesis comparisons. A. Comparison among the three different modularity hypotheses (see Figure 1) on the **G** matrix for each sex separately. B. Posterior distribution of the modularity signal for **D** and **Ω** discriminated by sex.

The comparison of the modularity signal for H_A_ on **D** and **Ω** showed different intensities of between-module integration as measured by Z_CR_ (Figure 3B). While **D** appears to behave in a similar way to **G**, with the whole distribution of Z_CR_ falling below the zero line for both sexes, **Ω** shows weaker modularity signal, with distributions that overlaps zero. This suggests that, while population divergence followed the genetic modularity pattern, the adaptive landscape was not aligned with those same patterns. The investigation of others modularity hypothesis (H_B_ and H_C_) for **D** and **Ω** showed distributions that overlap or were superior to zero, respectively, suggesting that evolution and selection did not fit those hypotheses either.

### Sex-biased selection of wing morphology

For the multivariate ***FSTq****–F*ST analysis of both sexes combined, the posterior distribution of the average eigenvalues of ***FSTq*** (*<u>λ</u>*) was higher than the simulated ones under drift, suggesting a pervasive action of directional selection in the among-population differentiation (Figure 4A). This was also true for individual axes of variation, especially the leading six axes (λ_1-6_), which showed no overlap between the 95% highest density intervals and the null-expectation. Inspection of the between-population divergence shows that males have diverged proportionally more than females (Figure 4B). On the analysis of the females and males separately, both sexes rejected the null-hypothesis of drift on the global test 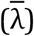, but only the leading axis of females had eigenvalues superior to the expected under drift (λ_1_), while males had at least three axes that could be reliably assigned to directional selection (λ_1-3_; Figure 5).

**Figure 4.**
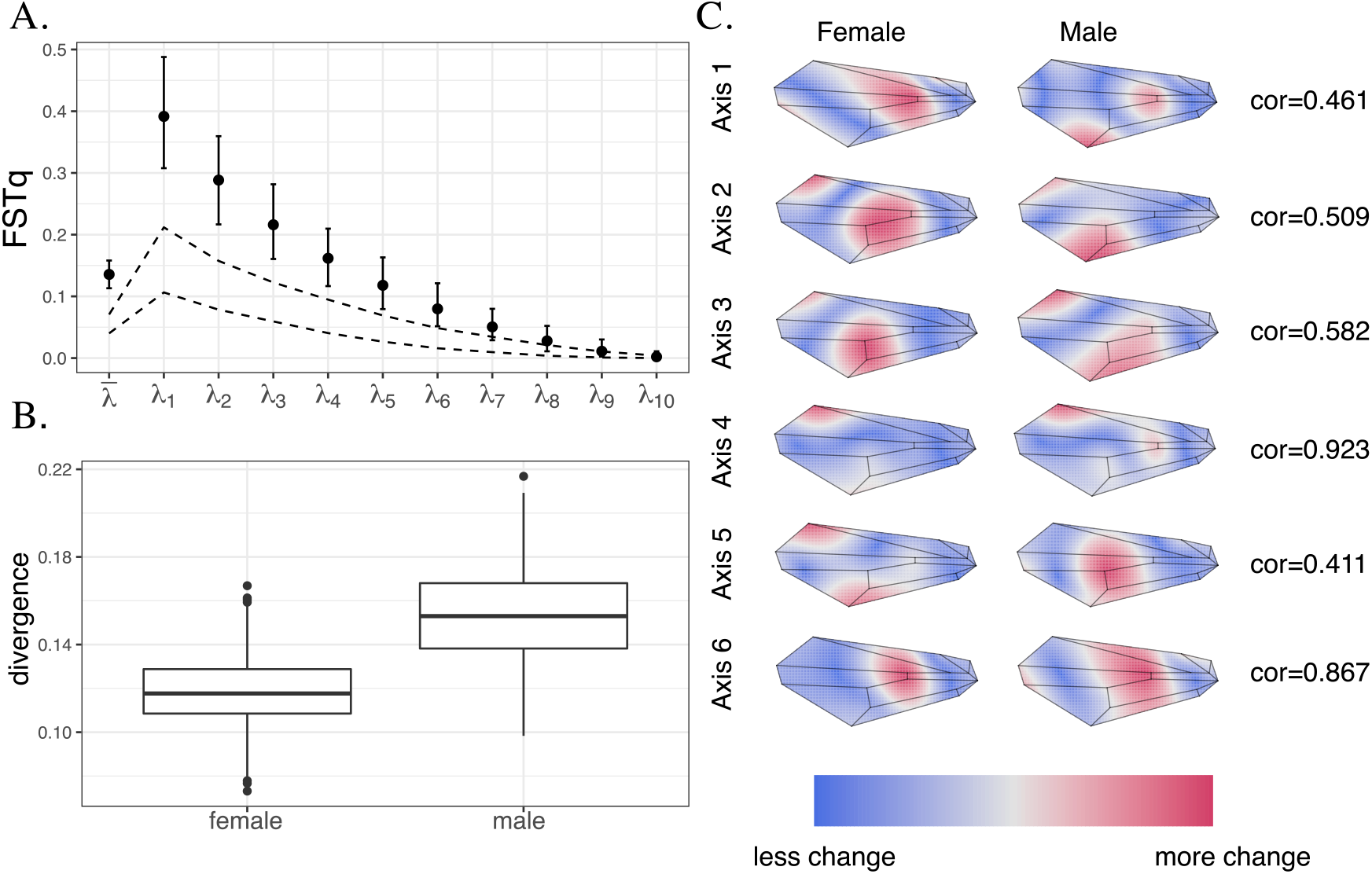
Tests of adaptive evolution using the eigenvalue decomposition of the *FSTq* matrix combining *D. buzzatii* male and female wing traits. **A**. Posterior distribution of eigenvalues of the ***FSTq*** matrix. Bars indicate posterior median and 95% highest density intervals for multivariate 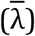 and univariate (λ_i_) genetic differentiation among populations. Dashed lines indicate upper and lower 95% highest density limits for the estimated ***FSTq*** under neutrality. **B**. Posterior distribution for the between-population amount of divergence. **C**. Representation of the leading six eigenvectors of the median ***FSTq*** matrix as shape deformations from the consensus configurations. Shapes are depicted as positive deviations from the consensus, and were amplified 20 times for visualization. Colors represent the amount of shape difference interpolated through a thin-plate spline. Lines represent the wing venation pattern. Correlation values are the vector correlation between female and male vectors.

An inspection of the shape changes associated with each axis of directional selection on the combined ***FSTq*** analysis reveal that males and females are evolving in different directions, a fact that is particularly true for the three leading eigenvectors, in which the correlation between the male and female-specific shape change are inferior to 0.58 (Figure 4C). This suggests that the leading axes of population differentiation represents a misalignment between phenotypic changes of males and females.

The *driftsel* analysis showed that, irrespective of the dataset (sexes combined or males and females separately), the resulting statistics **S** was always 1, reinforcing the pervasive role of directional selection shaping the wing-shape divergence.

### Effect of cross-sex (co)variances on the response to selection

The posterior distributions of **G** show that the intersexual correlations for the five PCs (r_mf_) were moderate to high. The 95% highest density intervals for r_mf_ values ranged from 0.502 to 0.763, suggesting a moderate to a high degree of between-sex pleiotropy on wing traits.

The R statistics based on the simulated random selection gradients shows a distribution that is centered on 1, and spreads evenly towards high and low values (Figure 6), suggesting that cross-sex covariances can either constrain or facilitate the evolution of *D. buzzattii* wing. On the other hand, the R values calculated based on empirically informed *β*s shows a distribution that is highly skewed to lower values (Figure 6), indicating that the observed populational differences occurred along directions that were in fact constrained.

**Figure 5.**
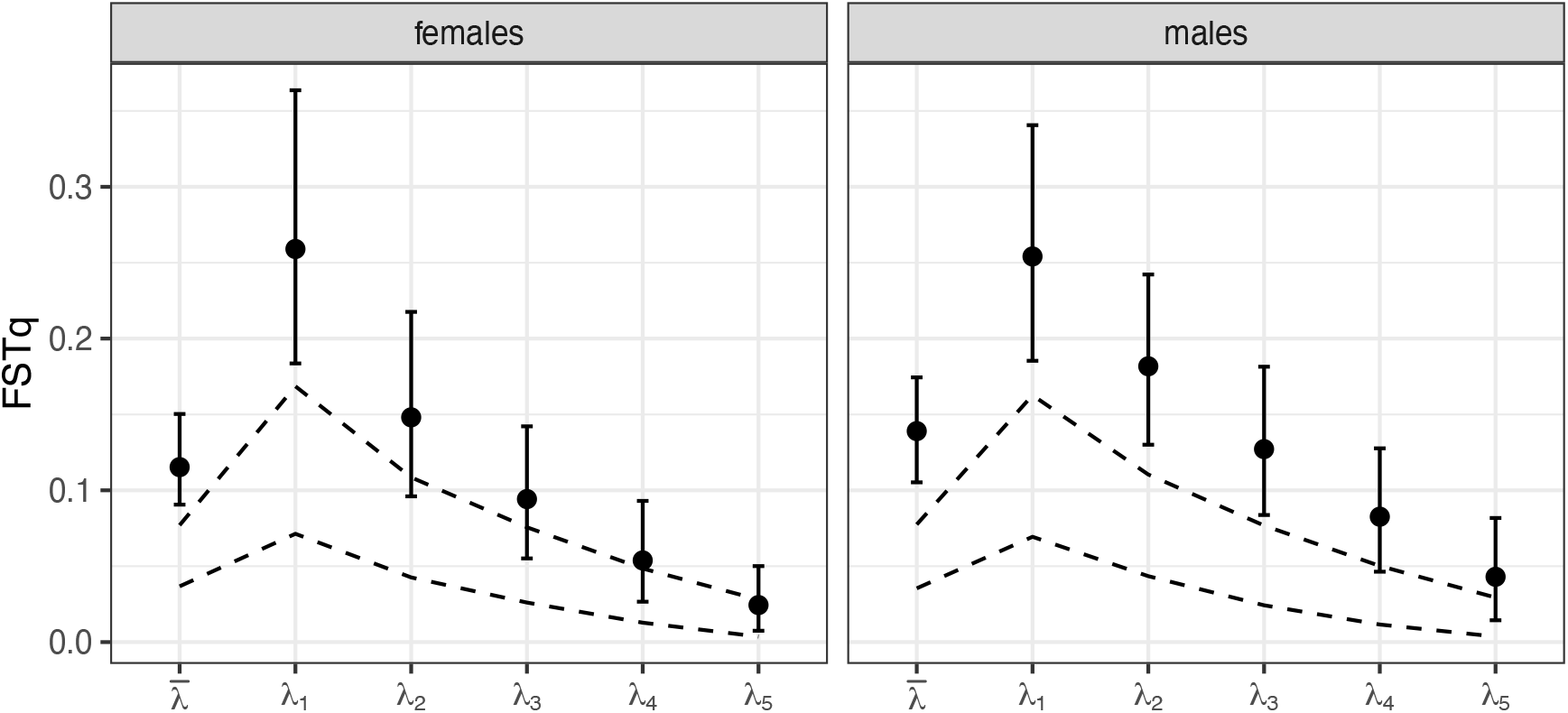
Tests of adaptive evolution for *D. buzzatii* wing traits using the eigenvalue decomposition of the *FSTq* matrix analyzing sexes separately. Bars indicate posterior median and 95% highest density intervals for multivariate 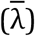 and univariate (λ_i_) genetic differentiation among populations. Dashed lines indicate upper and lower 95% highest density limits for the estimated ***FSTq*** under neutrality.

**Figure 6.**
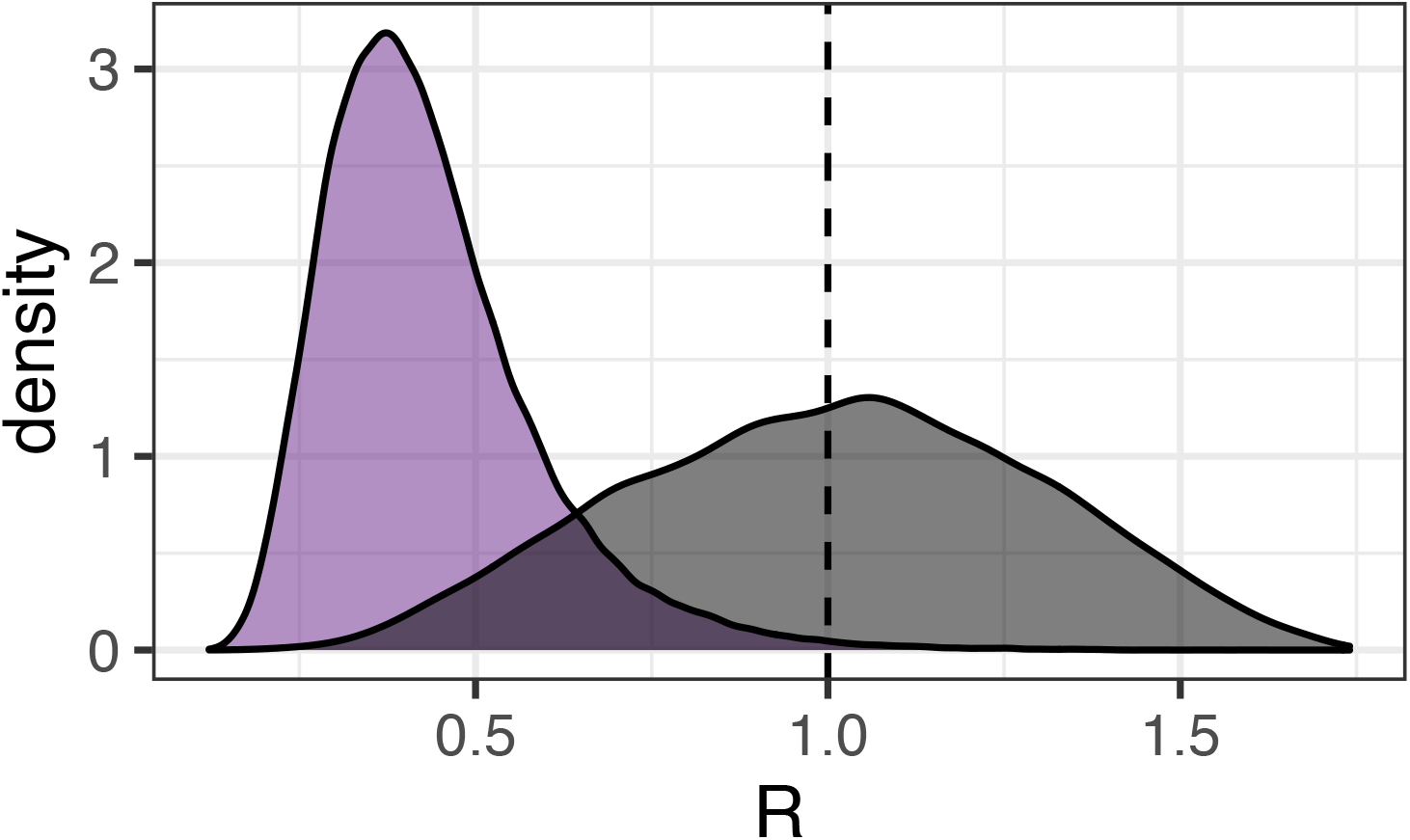
The R metric of random (gray distribution) and empirically informed (purple distribution) selection gradients. Dashed line indicates R=1, in which there is no constraining effect of between-sexes correlation.

## Discussion

We examined wing morphological changes in *D. buzzatii* in terms of constraints and evolutionary flexibility. We found that the wing blade and the wing base appear to be valid modules at the genetic level, providing developmental flexibility and phenotypic variation. However, reconstruction of between-population adaptive landscapes shows selection as an integrative force. Investigation of population’s morphological divergence suggests that each sex is diverging independently from each other, with males diverging more than females. As expected when sex-specific selection acts on a common genetic machinery (Bonduriansky & Chenoweth, 2009; Cox & Calsbeek, 2009), we found that populations are evolving in directions that are constrained by **G**, highlighting the restrictive role of between-sex pleiotropy in the evolution of sexually dimorphic traits.

Our multivariate QST analysis (***FSTq***) showed a clear signal of directional selection on wing shape, and that divergence was mostly concentrated on males rather than females. The leading eigenvectors of ***FSTq***, which are the axes of morphological variation most affected by directional selection, show moderate correlations between the directions of divergence between sexes, suggesting that males and females are evolving somewhat independently. Evaluations of between-population divergence show that greater divergence between sexes is usual. The ***FSTq*** analyses of each sex separately revealed three morphological axes under directional selection in males, but only one axis in females. This asymmetry suggests a wider array of functional demand on male wings than on female’s, which agrees with the double function of wings in males, like flight and courtship song production. Iglesias et al. (2018a) showed that males of these same sampled populations have divergent courtship songs, which they produce with their wings. These authors found evidence consistent with the role of directional selection in the divergence of courtship song traits, which could be associated with wing shape changes shown here. This male-specific selection on the wing for song production could also explain the asymmetry found in the number of axes under selection between male and female *D. buzzatii*. In summary, our results suggest that male wing shape is probably a target for multiple selective pressures, which lead to this sex diverging more than females in their phenotype.

The results of the evolutionary simulations showed that cross-sex covariances reduced the predicted response to selection in the direction of the extant sexual dimorphism, in line with previous results in *D. melanogaster’s* wing (Sztepanacz & Houle, 2019). However, while in *D. melanogaster* the predicted response to selection in random directions is also reduced, in *D. buzzatii* responses can be either reduced or augmented. This discrepancy may reflect the lower and more variable intersexual correlations (r_mf_) found in the wing of *D. buzzatii* (r_mf_ = 0.502-0.763), relative to values observed in *D. melanogaster’s* wing (r_mf_ = 0.907-0.940; Sztepanacz & Houle, 2019). Although lower than *D. melanogaster*, r_mf_ values found here were higher than, for instance, r_mf_ values found in the brightly colored dewlap of Anolis lizards (r_mf_ = 0.39–0.41; Cox, Costello, Camber, & McGlothlin, 2017). In *Anolis* lizards, cross-sex covariances (**B**) had little effect on the predicted response to selection either in random directions and in the direction of sexual dimorphism (Cox et al., 2017), reinforcing the idea that the intensity of correlations determines if populations will be constrained by **B**.

However, evaluation of between-population divergence of *D. buzzatii* shows that sexual dimorphism is greater when sex evolve in different directions, suggesting that selection along constrained directions had to be intensified in order to circumvent the limitations imposed by **G** (Machado, 2020), a fact that could have an indirect effect on the structure of **G** itself (Melo & Marroig, 2015). Theory shows that sexually antagonistic selection will favor a reduction in cross-sex genetic covariance when the strength of selection is highly asymmetric between the sexes (McGlodhlin et al., 2019). Otherwise, sexually antagonistic selection will tend to maintain strong cross-sex genetic covariance when the strength of selection is similar in each sex (McGlodhlin et al., 2019). If that holds for the populations investigated here, that means that the differences in r_mf_ values observed between *D. buzzatii* and *D. melanogaster* can be a consequence of different intensities and directions of sexual selection, with *D. buzzatii* being subjected to stronger sexual selection than *D. melanogaster*. Although both species rely on acoustic signals for mating success, the use of courtship song’s playbacks with wingless males does not reach the mating success of winged males in *D. melanogaster* (Rybak, Sureau, & Aubin, 2002), while they do in *D. buzzatii* (Iglesias & Hasson, 2017). This suggests that song preference is more relevant for the reproductive success of *D. buzzatii* males, and that the strength of selection related to female preference for courtship songs is stronger in this species than in *D. melanogaster*. Taken together, these results suggest that *D. buzzatii* is under stronger sexual antagonistic selection than *D. melanogaster*, leading not only to differences in patterns of divergence, but also having an impact on the genetic architecture of wing traits.

One possible genetic mechanism behind the decoupling of phenotypes among sexes could be related to the frequency of inversions in *D. buzzati*. A previous study in this species showed that inversion arrangements and the intensity of selection for adult viability on body size (a trait correlated with wing length and width; Norry, Vilarde, Fernandez Iriarte, & Hasson, 1997) are sex-specific (Rodriguez, Fanara, & Hasson, 1999). Changes in inversion frequencies after selection for adult viability (longevity) are concordant with expectations based on the average effect of inversions on body size and the intensity of selection on body size (Rodriguez et al. 1999). Moreover, changes in inversion frequencies are known to be under intense selection in *D. buzzatti* on a macrogeografic scale (Hasson, Rodriguez, Fanara, Reig, & Fontdevila, 1995; Rodriguez, Piccinali, Levi, & Hasson, 2000; Soto et al 2010). Given the differential frequency between the sexes, its relevance in trait determination, and the available evidence of natural selection, the patterns of variation in the inversion polymorphism may provide the genetic basis of sex-specific selection on morphological traits.

However, if selection was strong enough to overcome pleiotropic effects between sexes, why was it unable to change the modularity pattern of the wing into an integrated one, as expressed in the adaptive landscape? Studies of the association among traits in *D. melanogaster* wings suggests that, while it is possible to change trait integration through selection, these changes are not permanent and quickly revert to their ancestral patterns once selection is suspended (Bolstad et al., 2015). This suggests that internal stabilizing selection produced by the relation between ontogenetic pathways is an important component in maintaining integration patterns in complex traits, as it weeds away sub-optimal trait combinations due to maladaptive pleiotropic effects (Cheverud, 1988). However, changing between-sex integration does not necessarily result in (maladaptive) changes in the ontogenetic program, as phenotypes are still produced following the same general ontogenetic rules. If this argument holds, then it might explain why sexual dimorphism is more frequent than restructuration of integration patterns in complex phenotypes (Porto et al., 2009), even though both could be seen as instances of selection changing pleiotropic effects (Melo & Marroig, 2015).

While here we focused on the possible action of sexual selection in explaining between-sex divergence, it is possible that natural selection on wing shape, specifically through its relation to flight, could produce the observed differences as well. Wing shape is known to affect flight performance in *Drosophila* (Chin & Lentink, 2016; Ray, Nakata, Henningsson, & Bomphrey, 2016), and males are known to be the more dispersive sex in various species (Begon, 1976; Powell et al., 1976; Fontdevila & Carson, 1978; Markow & Castrezana, 2000; Mishra et al., 2020). Sex bias in dispersal is affected by many factors and interactions such as predispersal context, mate shortage, and availability of resources, which might vary among populations and exert different selective pressures (Mishra et al., 2018; Tung et al., 2018). Therefore, it is possible that natural selection for aerodynamics could produce between-sex coordinated and uncoordinated evolution. This is consistent with our ***FSTq*** analysis, which showed a mixture of aligned and non-aligned changes between sexes. Despite this, previous investigations into the evolution of sexually dimorphic traits in *Drosophila* have suggested that sexual selection has a stronger capacity to produce between-sex divergence (Chenoweth et al., 2008), suggesting that, even if aerodynamics affect the evolution of sexual dimorphism, it is more likely obfuscated by the role of sexual selection.

In short, the evolution of *D. buzzatii* wing shape seems to be the product of a complex interplay between ontogenetic constraints, conflicting sexual and natural selections, and presents a natural experiment for the evolution of sexual dimorphism on complex morphologies. Future studies on the causal links between wing morphology, song production and aerodynamics of *D. buzzatii* are needed to provide us with a better picture of how wings and cross-sex covariances can indirectly evolve as a by-product of selection on mating success and biomechanical performance.

## Acknowledgements

We sincerely thank Gladys Hermida for allowed access to its laboratory facilities for the production of the photographs used in this study. We also thank Consejo Nacional de Investigaciones Científicas y Técnicas (CONICET), ANPCyT (PICT 2013-1121 and PICT-2019-2554) and NSF (DEB 1350474 and DEB 1942717 to L. Revell) for financial support.

## Data Archiving Statement

The data that support the findings of this study will be openly available upon acceptance on Dryad (http://datadryad.org/) or Github (https://github.com/).

## Author Contributions

P.P.I., F.A.M. and E.M.S. conceived the project; E.H. supervised the global project in which the present work is embedded and contributed materials and reagents; E.M.S. performed wing dissections; S.L. took the photographs; P.P.I. and F.A.M. conducted analyses; P.P.I. and F.A.M. wrote the original draft, which was discussed, edited and revised by all authors.

